# Activation of extrasynaptic kainate receptors drives hilar mossy cell activity

**DOI:** 10.1101/2021.05.01.442200

**Authors:** Czarina Ramos, Stefano Lutzu, Miwako Yamasaki, Yuchio Yanagawa, Kenji Sakimura, Susumu Tomita, Masahiko Watanabe, Pablo E. Castillo

**Author notes:** To whom correspondence should be addressed: Pablo E. Castillo, MD/PhD, Dominick P. Purpura Department of Neuroscience, Albert Einstein College of Medicine, 1410 Pelham Parkway South, Kennedy Center, Room 703, Bronx, NY 10461, USA. These authors contributed equally to this work (Czarina Ramos, Stefano Lutzu). The authors declare no conflict of interest.

## Abstract

Mossy cells (MCs) of the dentate gyrus (DG) are key components of an excitatory associative circuit established by reciprocal connections with dentate granule cells (GCs). MCs are implicated in place field encoding, pattern separation and novelty detection, as well as in brain disorders such as temporal lobe epilepsy and depression. Despite their functional relevance, little is known about the determinants that control MC activity. Here, we examined whether MCs express functional kainate receptors (KARs), a subtype of glutamate receptors involved in neuronal development, synaptic transmission and epilepsy. Using mouse hippocampal slices, we found that bath application of submicromolar and micromolar concentrations of the KAR agonist kainic acid induced inward currents and robust MC firing. These effects were abolished in *GluK2* KO mice, indicating the presence of functional GluK2-containing KARs in MCs. In contrast to CA3 pyramidal cells, which are structurally and functionally similar to MCs, and express synaptic KARs at mossy fiber (MF) inputs (i.e., GC axons), we found no evidence for KAR-mediated transmission at MF-MC synapses, indicating that most KARs at MCs are extrasynaptic. Immunofluorescence and immunoelectron microscopy analyses confirmed the extrasynaptic localization of GluK2-containing KARs in MCs. Finally, blocking glutamate transporters, a manipulation that increases extracellular levels of endogenous glutamate, was sufficient to induce KAR-mediated inward currents in MCs, suggesting that MC-KARs can be activated by increases in ambient glutamate. Our findings provide the first direct evidence of functional extrasynaptic KARs at a critical excitatory neuron of the hippocampus.

**Significance Statement:** Hilar mossy cells (MCs) are an understudied population of hippocampal neurons that form an excitatory loop with dentate granule cells. MCs have been implicated in pattern separation, spatial navigation, and epilepsy. Despite their importance in hippocampal function and disease, little is known about how MC activity is recruited. Here, we show for the first time that MCs express extrasynaptic kainate receptors (KARs), a subtype of glutamate receptors critically involved in neuronal function and epilepsy. While we found no evidence for synaptic KARs in MCs, KAR activation induced strong action potential firing of MCs, raising the possibility that extracellular KARs regulate MC excitability *in vivo* and may also promote dentate gyrus hyperexcitability and epileptogenesis.

## Introduction

Hilar mossy cells (MCs) in the hilus of the dentate gyrus (DG) are major excitatory neurons that widely project onto dentate granule cells (GCs) to control their activity (Buckmaster and Schwartzkroin, 1994; Hashimotodani et al., 2017; Scharfman, 2018; Botterill et al., 2019). Within the DG, MCs form an associative network with GCs, in which MCs receive extensive convergent excitatory inputs from GCs (Patton and McNaughton, 1995; Acsady et al., 1998; Buckmaster and Jongen-Relo, 1999; Ribak and Shapiro, 2007) and then send excitatory feedback projections to up to ∼30,000 GCs along the dorsoventral axis of the ipsi- and contralateral hippocampus (Ribak et al., 1985; Frotscher et al., 1991; Buckmaster et al., 1996; Wenzel et al., 1997). Thus, the activity of a single MC can significantly impact the activity of numerous GCs, and ultimately, DG-CA3 information transfer. MCs contribute to hippocampal-dependent computations and behaviors such as pattern separation, spatial navigation and other cognitive functions such as novelty detection (Duffy et al., 2013; Danielson et al., 2017; GoodSmith et al., 2017; Senzai and Buzsaki, 2017; Fredes et al., 2021). In addition, aberrant MC function has been linked to brain disorders such as temporal lobe epilepsy (TLE), depression, anxiety and schizophrenia (Scharfman, 2016). Despite the important role that MCs play in brain function and disease, the mechanisms through which MC activity is recruited are still largely unexplored.

MCs share many structural and functional properties with CA3 pyramidal cells. Both CA3 pyramidal cells and MCs receive in their proximal dendrites a major excitatory input from GCs via the mossy fibers axons (MF), which impinge on complex spines called thorny excrescences (TEs) via giant presynaptic boutons (Amaral and Dent, 1981; Acsady et al., 1998). Functionally, the MF-to-MCs (MF-MC) synapse expresses robust forms of short- and long-term plasticity (Lysetskiy et al., 2005), similar to those reported at the MF-to-CA3 pyramidal cell (MF-CA3) synapse (Henze et al., 2002; Nicoll and Schmitz, 2005). In addition, glutamate release at both synapses is inhibited by activation of presynaptic group 2/3 metabotropic glutamate receptors (mGluR2/3) (Kamiya et al., 1996; Lysetskiy et al., 2005). While excitatory transmission is mainly mediated by AMPA and NMDA ionotropic glutamate receptors, a slow component of MF-CA3 synaptic transmission is mediated by kainate receptors (KARs) (Castillo et al., 1997). A similar, albeit modest KAR-mediated component has recently been reported at the MF-MC synapse (Hedrick et al., 2017). KARs are ionotropic glutamate receptors expressed in several brain areas, which have been implicated in neuronal development, neuronal excitability, synaptic transmission and plasticity

(Lerma and Marques, 2013). *In vivo* injection of the KAR agonist kainic acid (KA), a widely used animal model of TLE (Rusina et al., 2021), strongly activates CA3 pyramidal neurons (Westbrook and Lothman, 1983; Crepel and Mulle, 2015), and also leads to extensive MCs loss (Buckmaster and Jongen-Relo, 1999; Sloviter et al., 2003), suggesting that MCs are particularly sensitive to the activation of KARs. Intriguingly, while KAR-mediated responses are much weaker at MF-MC synapses (Hedrick et al., 2017) than at MF-CA3 synapses (Castillo et al., 1997; Mulle et al., 1998), transcriptome profiling revealed that MCs and CA3 pyramidal cells display comparable levels of transcripts for the functional KAR subunit GluK2 (Cembrowski et al., 2016), raising the possibility that KARs may have additional roles at MCs.

In this study, we combined *in vitro* electrophysiology in acute rat and mouse hippocampal slices, of wild type and *GluK2* knockout (KO) mice, with anatomical approaches such as immunofluorescence and immunoelectron microscopy, to determine the role and subcellular localization of KARs in MCs. We found that submicromolar and micromolar concentrations KA induced robust inward currents and strong MC firing, and both effects were absent in *GluK2* KO mice. Surprisingly, unlike in CA3 pyramidal neurons (Castillo et al., 1997), MF activation did not elicit any measurable KAR-mediated synaptic response in MCs. Consistent with these observations, immunofluorescence and immunoelectron microscopy revealed GluK2-containing KARs in the soma and dendrites of MCs, but nearly absent from MC TEs. Lastly, blocking glutamate uptake by excitatory amino-acid transporters (EAATs) elicited KAR-mediated inward currents in MCs. Altogether, our findings support the notion that MCs express functional extrasynaptic KARs whose activation by pharmacological agents (e.g., KA) and ambient glutamate may play an important role in engaging the GC-MC-GC recurrent circuit.

## METHODS

### Animals

Experiments were performed on postnatal Sprague-Dawley rats (P18-P28) of both sexes and C57BL/6 mice of both sexes for electrophysiological recordings. Animals were group-housed in a standard 12h light/12h dark cycle. WT, *GluK2* KO, and GAD67^+/GFP^ mice (Tamamaki et al., 2003); were obtained from Dr. Yanagawa, Gunma University, Japan. Handling and use of animals adhered to a protocol approved by the Animal Care and Use Committee at the Albert Einstein College of Medicine, at Yale University and at the Faculty of Medicine at Hokkaido University and in accordance with guidelines provided by the National Institutes of Health.

### Hippocampal Slice Preparation

Animals were deeply anesthetized with isoflurane and then decapitated. The brain was then rapidly removed from the skull and then hippocampi were dissected. Hippocampi were included in agar supports and acute transverse hippocampal slices (400 µm thick for Sprague-Dawley rats, 300 µm for C57BL/6 mice) were cut using a VT1200s vibratome (Leica Microsystems Co.) in a sucrose-based cutting solution containing (in mM): 215 sucrose, 2.5 KCl, 26 NaHCO_3_, 1.6 NaH_2_PO_4_, 1 CaCl_2_, 4 MgCl_2_, 4 MgSO_4_, and 20 D-glucose. After 15 mins in recovery post-sectioning, the solution was replaced by extracellular artificial cerebrospinal fluid (ACSF) recording solution containing (in mM): 124 NaCl, 2.5 KCl, 26 NaHCO_3_, 1 NaH_2_PO_4_, 2.5 CaCl_2_, 1.3 MgSO_4_, and 10 D-glucose. Slices were incubated for a minimum of 30 minutes in the ACSF before recording. Solutions were equilibrated with 95% O_2_ and 5% CO_2_ (pH 7.4).

### Electrophysiology

All experiments were performed in a submersion-type recording chamber perfused at ∼2 mL min^-1^, at 28 ± 1°C, except those in **Fig. 6** where temperature was raised to 34°C± 1°C. Whole-cell patch clamp recordings using a Multiclamp 700A amplifier (Molecular Devices) were performed from MCs and GCs in voltage-clamp (V_hold_ −60 mV), or in current-clamp configuration (V_rest_ ∼-65 mV) using borosilicate pipette electrodes (∼3-4 MΩ). Recordings were performed using a K^+^-based internal solution containing (in mM): 135 KMeSO_4_, 5 KCl, 1 CaCl_2_, 5 NaOH, 10 HEPES, 5 MgATP, 0.4 Na_3_GTP, 5 EGTA, 10 D-glucose, pH 7.2 (280-290 mOsm). In some recordings we also employed a Cs^+^-based internal solution containing (in mM): 131 Cs-gluconate, 8 NaCl, 1 CaCl_2_, 10 EGTA, 10 D-glucose and 10 HEPES, pH 7.2 (285-290 mOsm). Series resistance (∼7-25 MΩ) was monitored throughout all experiments with a −5 mV, 80 ms voltage step, and cells that exhibited a significant change (>20%) were excluded from analysis.

MCs were identified using previously established criteria (Larimer and Strowbridge, 2008). Specifically, we measured firing properties and membrane time constant by injection of a step of depolarizing current while in current-clamp configuration. Cells were confirmed as MCs by exhibiting elevated spontaneous synaptic activity, little to no afterhyperpolarization and non-burst firing patterns upon depolarizing pulses (5s duration, 60-120 pA). Additional confirmation was performed post-hoc through morphological analysis of biocytin-filled cells where MCs were identified by the presence of distinctive complex TEs in their proximal dendrites. To isolate KAR-mediated currents and EPSCs, we bath applied a cocktail of antagonists which included: LY303070 (15 µM) or GYKI 53655 (GYKI – 30 µM), D-APV (25 µM), picrotoxin (50 µM) and CGP35348 (3 µM) to block AMPA, NMDA, GABA_A_ and GABA_B_ receptors, respectively. The cocktail was applied right after the target cell was identified as a MC. For the isolation of KAR-EPSC, we first monitored AMPAR-EPSCs in presence of the above specified cocktail, without LY303070, which was bath applied after a stable baseline was acquired.

To evoke MF synaptic responses in MCs and CA3 pyramidal cells, a bipolar stimulating theta-glass pipette was filled with ACSF and placed in the subgranular zone of the DG. Only EPSCs that showed < 2 ms 20-80% rise time and robust paired-pulse facilitation (EPSC2/EPSC1 > 2) were considered MF-derived and included in the analysis.

To increase the probability of detecting a KAR-mediated EPSC, we delivered two stimuli (5 ms inter-stimulus interval, 100 µs duration, ∼100 µA amplitude), using a stimulus isolator unit (Isoflex, AMPI). Typically, stimulation was adjusted to obtain comparable magnitude synaptic responses across experiments.

### Data analysis for electrophysiology experiments

Electrophysiological data were acquired at 5 kHz filtered at 2.4 kHz and analyzed using custom-made software for IgorPro (Wavemetrics Inc.). The change in the holding current (ΔI holding) was calculated by subtracting the baseline holding current value (average of 50 s before KA application) from the average holding current post-drug application (average 50 s before washout). To calculate firing rate in the current clamp configuration, spikes were detected using a custom-made MatLab script, which detected all voltage increases above a threshold value established by the experimenter (i.e., 0 mV). When 3 µM KA was used to depolarize MCs, the peak of the action potentials gradually decreased (likely due to inactivation of voltage-gated sodium channels) and became undistinguishable from spontaneous activity. This led to an underestimation of the effect of 3 µM KA on MCs firing. Firing rate was quantified as number of spikes per second.

### Reagents

Reagents were bath applied following dilution into ACSF from stock solutions stored at −20°C prepared in water or DMSO, depending on the manufacturer’s recommendation. The final DMSO concentration was <0.01% total volume. All chemicals and drugs used for the electrophysiology experiments were purchased from Sigma-Aldrich (St. Louis, MO, USA) except NBQX, CGP-55845, DCG-IV, and GYKI 53655, which were obtained from Tocris-Cookson (Minneapolis, MN, USA), and tetrodotoxin, which was obtained from HelloBio (Inc, Princeton, NJ). LY 303070 was obtained from ABX advanced biochemical compounds (Radeberg, Germany).

### Quantification and statistical analysis

Statistical analysis was performed using OriginPro software (OriginLab). The normality of distributions was assessed using the Shapiro-Wilk test. In normal distributions, Student’s unpaired and paired t Tests were used to assess between-group and within-group differences, respectively. The non-parametric paired sample Wilcoxon signed rank test and Mann-Whitney’s U test were used in non-normal distributions. Statistical significance was set to p < 0.05 (*** indicates p < 0.001, ** indications p < 0.01, and * indicates p < 0.05). All values are reported as the mean ± SEM.

### Fixation and sections

We used glyoxal fixative containing 9% glyoxal and 8% acetic acid (v/v, pH 4.0 adjusted with 5N NaOH), which is modified from the original glyoxal fixative (Richter et al., 2018). Under deep pentobarbital anesthesia (100 mg/kg body weight, i.p.), mice were fixed by transcardial perfusion with ∼60 ml of glyoxal solution for 10 min at room temperature. Brains were postfixed in the same fixative for 3 h and cryoprotected with 30% sucrose in 0.1 M PB (pH 7.2) for 2 d. For immunofluorescence and immunoelectron microscopy, 50-μm-thick coronal sections through the ventral hippocampus (3.0–3.7 mm posterior to Bregma) were prepared on a cryostat (CM1900; Leica Microsystems) and subjected to free-floating incubation.

### Antibodies

We used the following antibodies: mouse anti-calretinin (MAB1568, Millipore; RRID, AB_94259), goat anti-EGFP (Takasaki et al., 2010)(AB_2571574), rabbit anti-GluK2/3 (Straub et al., 2011), guinea pig anti-Neto1(Straub et al., 2011), and guinea pig anti-PSD95 (Fukaya and Watanabe, 2000)(AB_2571612).

### Immunofluorescence

All immunohistochemical procedures for immunofluorescence were performed at room temperature and PBS containing 0.1% Triton-X100 was used as a dilution and washing buffer. Sections were incubated with 10% normal donkey serum for 20 min, a mixture of primary antibodies overnight (1 μg/ml each), and a mixture of Alexa Fluor 405-, 488-, 647-, or Cy3-labeled species-specific secondary antibodies for 2 h at a dilution of 1:200 (Invitrogen; Jackson ImmunoResearch). To avoid cross talk between multiple fluorophores, images were taken with a confocal laser-scanning microscope equipped with 405-, 473-, 559-, and 647-nm diode laser lines, and UPLSAPO 10× (NA, 0.4), and PLAPON 60×OSC2 (NA, 1.4; oil immersion) objective lenses (FV1200, Olympus). Image and pinhole size were 800 × 800 pixels and 1 airy unit, respectively. To compare genotypic and regional difference, images were taken at the same condition.

### Preembedding immunoelectron microscopy

All incubations were performed at room temperature and PBS containing 0.1% Tween20 was used as a dilution and washing buffer. Sections were incubated in 10% normal goat serum (Nichirei, Tokyo, Japan) for 20 min, and with primary antibody against GluK2/3 (1 µg/ml) overnight and then with secondary antibodies linked to 1.4-nm gold particles (1:100; Nanogold; Nanoprobes) for 4 h. After extensive washing with PBST and HEPES buffer (200 mM sucrose, 50 mM HEPES, pH 8.0), immunogold was intensified with a silver enhancement kit (R-GENT SE-EM; Aurion) for 45–60 min. Sections were further treated with 1% osmium tetroxide for 15 min, stained with 2% uranyl acetate for 20 min, dehydrated with graded ethanol series, and embedded in Epon 812 (TAAB). After polymerization at 60°C for 48 h, ultrathin sections were prepared with an ultramicrotome (Ultracut; Leica), mounted on copper-mesh grids and stained with 2% uranyl acetate for 5 min and Reynold’s lead citrate solution for 1 min. Photographs were taken with a JEM1400 electron microscope (JEOL, Tokyo, Japan). Electron micrographs were randomly taken within ∼5 µm from the surface to avoid false-negative areas. To quantify metal particle labeling, 3 × 3 montage images (∼6 µm × 6 µm) were randomly taken at a magnification of 15,000×.

For quantitative analysis, plasma membrane-attached immunogold particles, being defined as those apart <35 nm from the cell membrane, were counted and analyzed using MetaMorph software (Molecular Devices). The mean number of membrane-attached gold particles per 1 μm of the plasma membrane was counted for each neuronal compartment (dendritic spine, dendritic shaft, and soma). Measurements were made from three WT and two *GluK2* KO mice and pooled together, because there was no significant difference in the labeling density in the same genotype. In each neuronal compartment, labeling density was calculated for individual profile. Statistical analyses were performed using GraphPad Prism 9.0 software (GraphPad Software). All data are given as mean ± SEM. Data were analyzed using Kruskal-Wallis test followed by Dunn’s post Test. *p < 0.05; **p < 0.01; ***p < 0.001.

## Results

### Kainate receptors mediate inward currents and action potential firing in hilar mossy cells

To test whether MCs expressed functional KARs, we first performed whole-cell patch clamp recordings from MCs in acute rat hippocampal slices and bath applied the KAR agonist KA. MCs were identified based on the high frequency of spontaneous EPSCs, non-burst firing pattern upon depolarization, and action potentials with almost no afterhyperpolarization (see Methods) (Larimer and Strowbridge, 2008) **(Fig. 1A)**. To confirm the identity of the recorded cell, we loaded putative MCs with biocytin and stained with Alexa 594-conjugated streptavidin, and confirmed the presence of TEs, a hallmark of MCs (**Fig. 1A**) (Scharfman and Schwartzkroin, 1988). We examined whether KA bath application induced KAR-mediated inward currents in MCs, as previously shown in KAR-expressing CA3 pyramidal cells (Castillo et al., 1997; Mulle et al., 1998) To isolate these currents, recordings were performed in the presence of LY303070 (15 µM), D-APV (25 µM), picrotoxin (50 µM) and CGP35348 (3 µM) to block AMPARs, NMDARs, GABA_A_ and GABA_B_ receptors, respectively, and MCs were voltage clamped at −60 mV. Under these recording conditions, KA bath application (0.3 µM and 3 µM) induced large, concentration-dependent inward currents in MCs (**Fig. 1B,C**) (ΔI holding MC: 0.3 µM KA: 165.3 ± 28.56 pA; n = 5a/5c; 3 µM KA: 736.24 ± 82.34 pA; n = 4a/4c – one of the cells died after 0.3 µM application). In contrast, the same concentrations of KA induced modest currents in GCs (ΔI holding GC: 0.3 µM KA: 30.73 ± 1.38 pA; n = 3a/3c; 3 µM KA: 72.21 ± 6.31 pA; n = 3a/3c). By activating KARs in CA3 pyramidal cells, KA application could recruit CA3 pyramidal cells (Robinson and Deadwyler, 1981; Westbrook and Lothman, 1983; Castillo et al., 1997) which make synaptic contacts with MCs (Scharfman, 1994) and could indirectly activate KARs in MCs. To test this possibility, action potential generation was prevented by perfusing the voltage-gated sodium channel blocker tetrodotoxin (TTX, 0.5 µM) in the bath. In the presence of TTX, KA-induced currents were not significantly different from control conditions (**Fig. 1D,E**) (ΔI holding MC + TTX, 0.3 µM KA : 117.3 ± pA; n = 5a/6c; 3 µM KA: 765.19 ± 79.06 pA; n = 5a/6c; 0.3 µM control vs TTX: n.s. p = 0.12045, two sample t Test; 3 µM control vs TTX: n.s. p = 0.81287: two sample t Test), indicating that these currents do not result from indirect activation of CA3 pyramidal neurons. In addition, the competitive AMPAR/KAR antagonist NBQX (25 µM) abolished KA-mediated inward currents, strongly suggesting these currents were mediated by KAR activation in MCs (**Fig. 1D,E**) (ΔI holding MC 0.3 µM KA control: 182.78 ± 25.7 pA; n = 3a/4c; 3 µM KA + 25 µM NBQX: 12.99 ± 3.31; n = 3a/4c; 0.3 µM KA vs 3 µM KA + NBQX: *** p = 0.000012: two sample t Test). Lastly, KA bath application induced currents in MCs of mouse hippocampal slices,which were abolished in *GluK2* KO mice (**Fig. 1F,G**) (ΔI holding MC WT; 0.1 µM KA: 58.33 ± 19.49 pA, 0.3 µM KA: 75.44 ± 13.87, 1 µM KA: 233.91 ± 15.67 pA, 3 µM KA: 494.32 ± 85.01 pA; ΔI holding *GluK2* KO; 0.1 µM KA: 19.77 ± 9.27 pA, 0.3 µM KA: 13.35 ± 4.44 pA, 1 µM KA: 24.74 ± 10.55 pA, 3 µM KA:14.39 ± 5.04 pA; WT vs *GluK2* KO: *F*(1,3) = 146.98864, ** p = 0.00121; two-way ANOVA repeated measures), indicating that these currents are mediated by GluK2-containing KARs.

**Figure 1.**
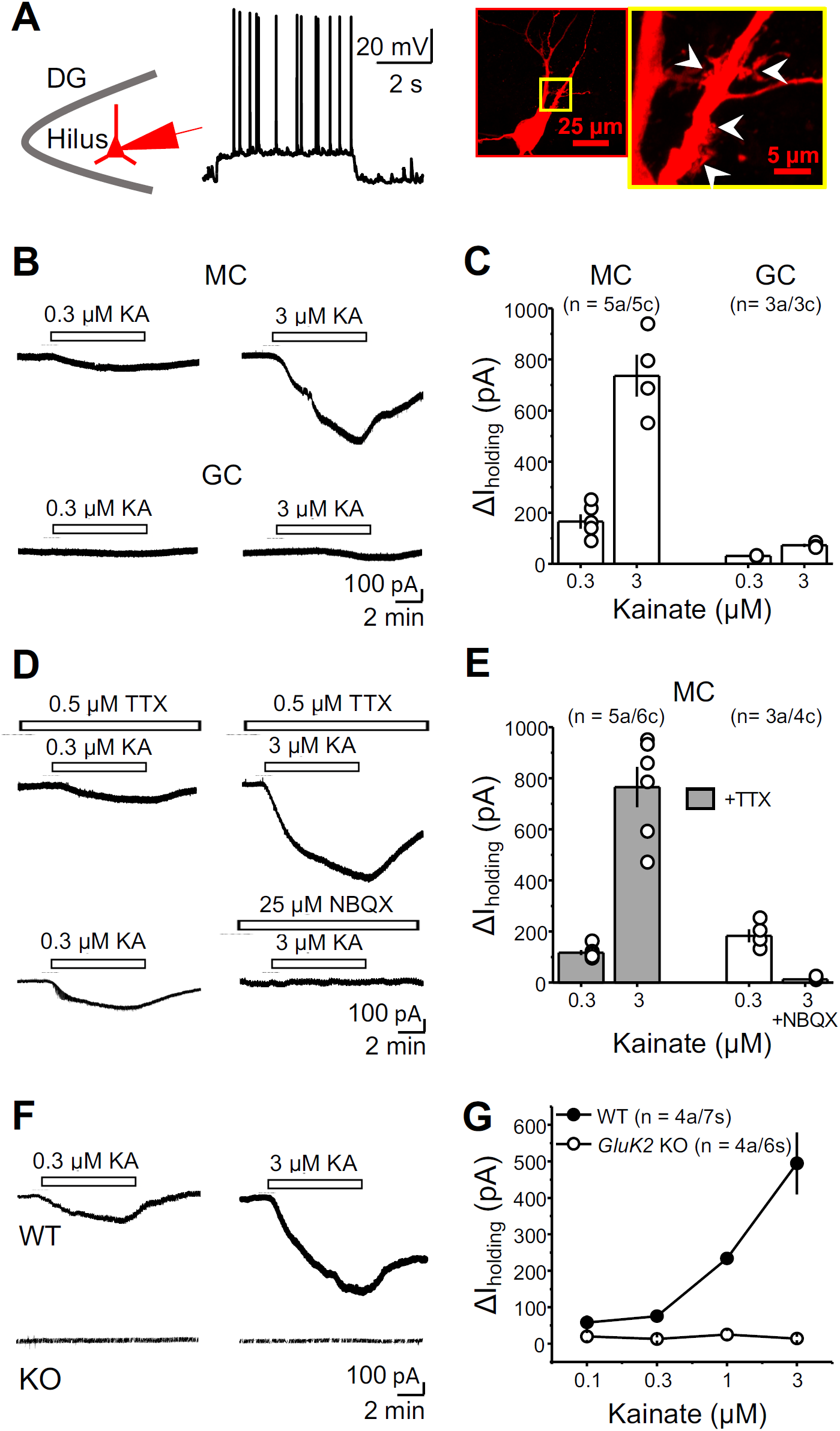
Activation of KARs mediates inward currents in hilar mossy cells. (**A**) Identification of hilar mossy cells (MCs) in acute hippocampal slices. *Left*, schematic of MC recordings in the hilus of the dentate gyrus (DG). *Middle*, patched neurons were depolarized to analyze their firing properties. *Right*, post-hoc staining of a MC using Alexa 594-conjugated streptavidin. The yellow boxed area is magnified on the right-hand side. Patched cells were confirmed to be MCs if evoked spikes displayed no evident afterhyperpolarization and by the presence of thorny excrescences (TEs) (white arrowheads). (**B**) Representative experiment showing that bath application of 0.3 and 3 µM kainic acid (KA) in the same cell induced concentration-dependent inward currents in MCs (*top*), but only a negligible inward current in dentate gyrus granule cells (GC) (*bottom*). (**C**) Summary plot of the amplitude of the inward currents (ΔI holding) induced by KA application. (**D**) Representative experiments showing that KA-induced inward current was not affected by co-application of TTX (0.5 µM) (*top*) but it was abolished by the AMPAR/KARs antagonist NBQX (25 µM) (*bottom*). A low concentration of KA (0.3 µM) was previously tested to verify the presence of a normal, fully reversible KA-induced current in the same cell. (**E**) Summary plot. (**F**) KA-induced currents in MCs were robust in WT (*top*) but abolished in *GluK2* KO mice (*bottom*). (**G**) Concentration-response curve in WT and *GluK2* KO mice. Data are presented as mean ± S.E.M.

We hypothesized that KAR-mediated currents can produce enough depolarization to drive MC action potential firing. To test this possibility, we recorded MCs in current-clamp mode before and after KA application. Given that hippocampal interneurons impinging on MCs could express functional KARs (Frerking et al., 1998), these experiments were performed with intact excitatory and inhibitory components of synaptic transmission in order to assess the net effect of KAR activation on MC firing. Under these recording conditions, bath application of 0.3 µM and 3.0 µM KA induced strong MC firing (**Fig. 2A**) (WT average firing rate 0.3 µM KA: 0.333 µM ± 0.025 spikes/s; n = 2a/3c; 3 µM KA: 5.54 ± 0.28; n = 2a/3c), and this effect was abolished in *GluK2* KO mice (**Fig. 2B**). These results indicate that activation of GuK2-containing KARs with low concentrations of the agonist KA can powerfully drive MCs.

**Figure 2.**
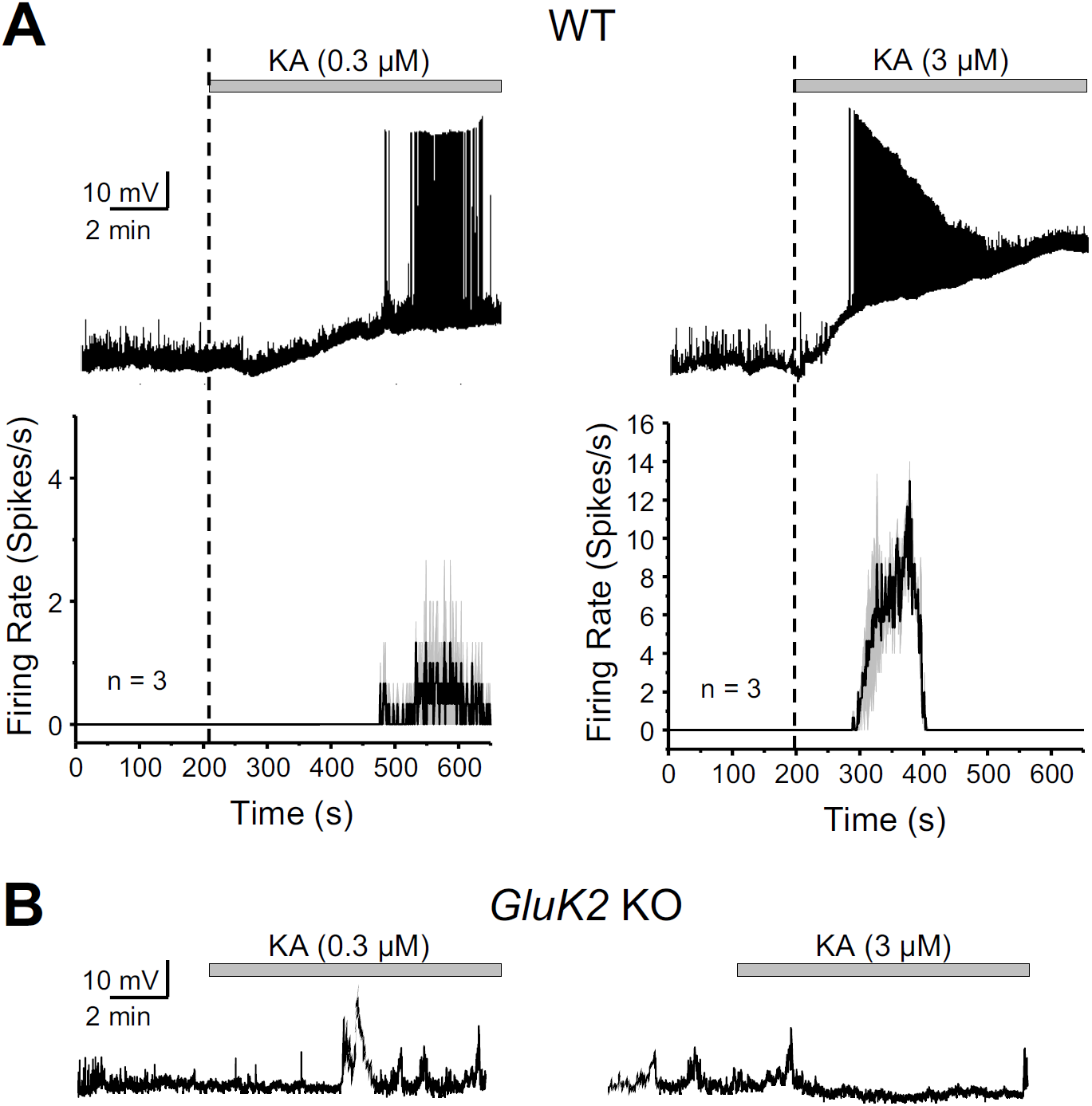
GluK2-containing KARs mediate KA-induced robust action potential firing of MCs. (**A**) *Top*, Effect of of KA bath application (0.3 and 3 µM) on MCs membrane potential in current-clamp configuration. 3 µM KA application induced firing with faster onset and higher frequency than 0.3 µM KA. Firing in 3 µM KA eventually disappeared likely due to excessive depolarization and action potential refractoriness. *Bottom*, average firing rate histogram (expressed as spikes per second) for the experiments in top panel. Black and grey traces represent mean and S.E.M. of MCs firing rate. (**B**) Representative traces showing no effect of KA application on MCs firing in *GluK2* KO mice.

### KARs-mediated EPSCs are undetectable at MF-MC synapse

We next sought to determine the subcellular localization of KARs on MCs. Because of the strong structural and functional similarities between CA3 pyramidal cells and MCs, we tested whether MF activation elicits KAR-EPSCs in MCs as previously shown at MF inputs onto CA3 pyramidal cells (Castillo et al., 1997). To this end, we evoked AMPAR-EPSCs by stimulating MF axons with two stimuli to boost glutamate release from MFs (5 ms inter-stimulus interval) in the presence of a cocktail of NMDARs, GABA_A_ and GABA_B_ receptor antagonists (see Methods). We then attempted to isolate the KAR-mediated component of the MF-EPSC by applying the selective, non-competitive AMPAR antagonist GYKI 53655 (30 µM). GYKI application abolished the AMPAR-EPSCs but surprisingly, it failed to uncover a KAR-mediated EPSC (**Fig. 3A**) (EPSC amplitude post GYKI application: 2.35 ± 0.52 % of baseline; n = 4a/5s). In addition, increasing the number of stimuli (from 2 to 5), a manipulation expected to increase the likelihood of detecting KAR-EPSCs at the MF-CA3 synapse (Castillo et al., 1997), did not generate any detectable current either (data not shown; 5 pulses: 2.108 ± 0.415 % of baseline; n = 4a/5s). To verify that the EPSCs were MF-mediated, in a separate set of experiments, we applied the mGluR2/3 agonist DCG-IV (1 µM) which selectively blocks glutamate release at MF-MC synapses (**Fig. 3B**) (Lysetskiy et al., 2005; Hedrick et al., 2017). DCG-IV reduced synaptic responses by ∼70 %, indicating our stimulation mainly recruited MF inputs onto MCs (EPSC amplitude post DCG IV: 29.35 ± 7.07 % of baseline; n = 3a/4s). As a positive control, and as previously reported (Castillo et al., 1997), MF stimulation elicited GYKI-resistant, NBQX-sensitive KAR-EPSCs in CA3 pyramidal neurons (**Fig. 3C**) (EPSC amplitude post GYKI application: 15.85 ± 6.59 % of baseline; n = 2a/4s; KAR-EPSC amplitude post NBQX application: 0.47 ± 0.25 % of baseline; n = 2a/4s). These results indicate that in contrast to MF-CA3 synapses, KARs do not mediate synaptic transmission at MF-MC synapses. Given the robust activation of MCs by low concentrations of KA (Figs. 1,2), our findings thus far suggest KARs in MCs are extrasynaptic.

**Figure 3.**
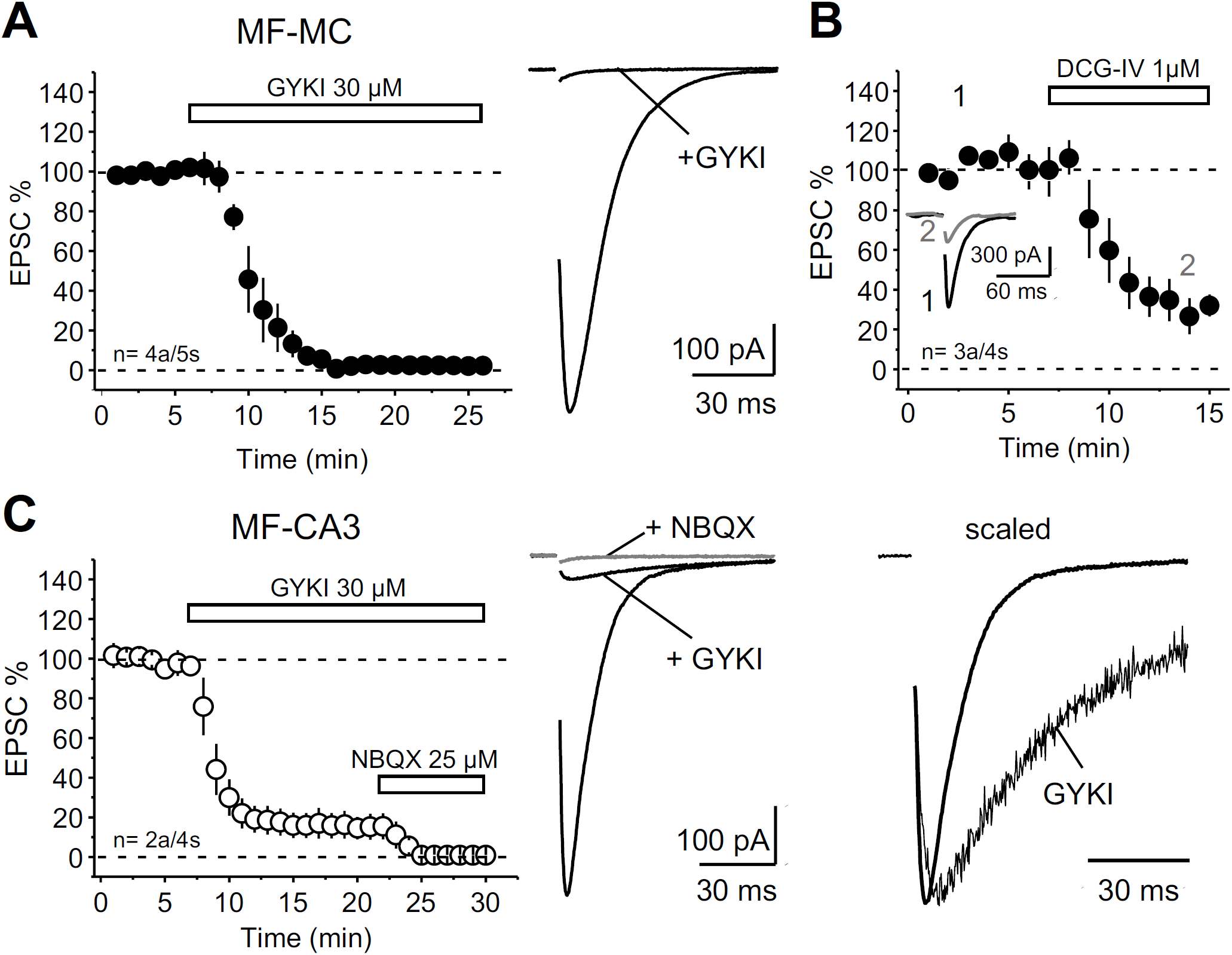
KAR-mediated transmission at MF-CA3 but not MF-MC synapses. (**A**). *Left*, Bath application of the selective AMPAR antagonist GYKI 53655 (30 µM) abolished synaptic transmission at MF-MC synapses. *Right*, representative average traces (30 consecutive responses). (**B**) Bath application of the mGluR2/3 agonist DCG-IV (1 µM) significantly reduced MF-MC EPSCs (∼70%). (**C**) *Left*, GYKI 53655 application (30 µM) blocked AMPAR-mediated transmission at the MF-CA3 synapse, thus revealing a KAR-mediated component that was blocked by 25 µM NBQX. *Middle*, representative average traces (30 consecutive responses) before and after GYKI application, and after subsequent application of NBQX. *Right*, normalized AMPAR and KAR components highlighting the slow kinetics of the KAR-EPSC. In all traces included in this figure, stimulus artifacts were deleted for clarity. Data are presented as mean ± S.E.M.

### Distinct subcellular localization of KARs in CA3 pyramidal cells and mossy cells

To determine the anatomical localization of KARs in MCs, we applied immunostaining to tissue sections fixed with a glyoxal-based fixative (see Methods), which is effective for detection of both non-synaptic and synaptic molecules. First, we confirmed the specificity of the antibody against GluK2/3 by blank labeling in *GluK2* KO hippocampus (**Fig. 4A,B**). In WT mice, the antibody yielded a contrasting pattern of labeling across hippocampal subregions: intense and coarse punctate labeling in the CA3 *stratum lucidum*, and moderate and diffuse labeling in the hilus (**Fig 4A**). Further quadplex immunofluorescence using GAD67^+/GFP^ mice (Tamamaki et al., 2003) allowed us to distinguish excitatory MCs from inhibitory calretinin-positive interneurons, and to examine if MCs express GluK2/3 (**Fig. 4C,D**) together with or without the excitatory postsynaptic marker PSD95 (**Fig. 4C**), or the KAR auxiliary subunit Neto1 (**Fig. 4D**). In CA3 *stratum lucidum*, GluK2/3-positive puncta were intense and aggregated into large clusters and were almost perfectly overlapped with PSD95 (**Fig. 4E,F**), suggesting exclusive localization in MF-CA3 pyramidal cell synapse. Compared with CA3, GluK2/3 labeling was smaller and less frequent in the hilus of the DG (**Fig. 4G,H**). MC soma and dendrites, which can be unequivocally identified as calretinin-positive and GFP-negative structures (**Fig. 4G**), were associated with GluK2/3 puncta that did not colocalize with PSD95 (**Fig. 4H**). Similarly, while Neto1 staining was overlapped with GluK2/3-positive puncta in *stratum lucidum* (**Fig. 4I,J**), it was not found around MC soma and dendrites (**Fig. 4K,L**). Together, these findings suggest that GluK2/3-containing KARs in MCs are expressed at extrasynaptic sites.

**Figure 4.**
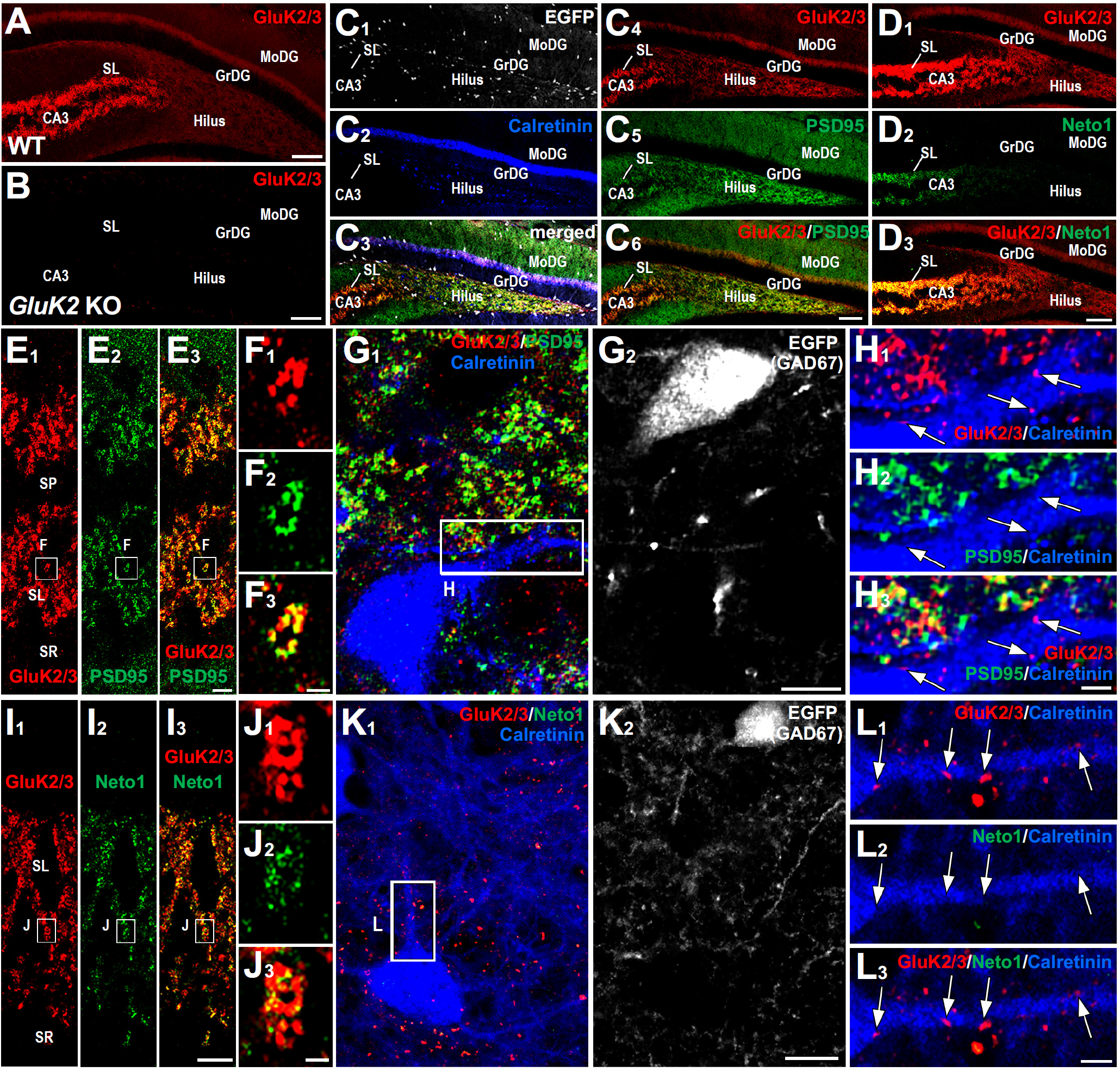
Contrasting localization of GluK2/3 and its molecular partners in CA3 *stratum lucidum* and DG hilus. (**A,B**) In WT mice, GluK2/3 labeling is intense in CA3 *stratum lucidum*, while it is moderate and diffuse in the dentate gyrus (A). Note the lack of GluK2/3 staining in *GluK2* KO mice, indicating the specificity of the GluK2/3 antibody and exclusive expression of GluK2 in these hippocampal regions (B). **(C)** Quadplex immunofluorescence for GFP (C_1_, white), calretinin (C_2,_ blue), GluK2/3 (C_4_, red), and PSD95 (C_5_, green) in GAD67^+/GFP^ mice. (**D**) Double immunofluorescence for GluK2/3 (D_1_, red) and Neto1 (D_2_, green). Note that intense signal for Neto 1 is almost limited to CA3 *stratum lucidum*. (**E-H**) Distinct GluK2/3 and PSD95 localization between CA3 pyramidal cells and hilar MCs. (**E,F**) GluK2/3 and PSD95 labeling are intense in the CA3 *stratum lucidum* (E). A high-magnification image confirms their extensive overlap (F). (**G,H**) A MC, which is identified as a calretinin-positive (G_1,_ blue) and GFP-negative (G_2,_ white) cell in GAD67^+/GFP^ mice, shows weak labeling for GluK2/3 on its dendrites (red, arrows in H). Note that such GluK2/3 puncta are neither overlapped nor associated with PSD95 signal (H_2_, green). (**I-L**) Distinct GluK2/3 and Neto1 localization between CA3 pyramidal cells and hilar MCs. (**I,J**) Intense GluK2/3 and Neto1 labeling in the CA3 *stratum lucidum* (l). A high-magnification image shows Neto1 labeling is only observed on GluK2/3-positive puncta (J). (**K,L**) A MC shows weak labeling for GluK2/3 on its dendrites (red, arrows in L) but lacks Neto1 labeling (L_2_, green). DG, dentate gyrus; GrDG, granule cell layer of the DG; MoDG, molecular layer of the DG; SL, *stratum lucidum*; SP, *stratum pyramidale*; SR, *stratum radiatum*. Scale bars, (**A,D**) 100 µm; (**E,F,I,J,G,K**) 10 µm; (**H,L**) 2 µm.

For a more accurate assessment of the subcellular localization of KARs in MCs, we performed pre-embedding immunoelectron microscopy for GluK2/3. In WT mice, metal particles for GluK2/3 were observed on the postsynaptic membrane of TEs (**Fig. 5A,G**, blue) of CA3 pyramidal cells facing large MF boutons (12.1 ± 0.95 particles/mm). In *GluK2* KO mice, immunolabeling was essentially absent on the postsynaptic membrane of CA3 spines (**Fig. 5B,G** blue), confirming the specificity of the immunolabeling (0.07 ± 0.07 particles/mm; Dunn’s multiple comparison test; *p* < 0.0001, compared to WT). In the DG hilus, MCs can be identified as having spiny dendrites contacted with large MF terminals (Acsady et al., 1998). In contrast to MF-CA3 synapses, MF-MC synapses were not labeled for GluK2/3 (**Fig. 5C,G**) (0.06 ± 0.04 particles/μm in WT, vs 0.01 ± 0.01 particles/μm in *GluK2* KO; *p* > 0.99). Instead, occasional weak labeling was observed on the non-synaptic membrane of dendritic shaft and spines of MCs (**Fig. 5E,F** green). The density of non-synaptic particles was much lower than that at MF-CA3 synapses (p = 0.0137), but significantly higher than their counterparts in *GluK2* KO (**Fig. 5D,G**; 4.1 ± 0.8 particles/μm in WT vs 0.02 ± 0.01 particles/μm in *GluK2* KO; *p* = 0.008). These results not only demonstrate the presence of KARs in MCs but also show that, in contrast to CA3 pyramidal neurons, GluK2-containing KARs are exclusively expressed at non-synaptic sites in proximity of MF-MC synapses.

**Figure 5.**
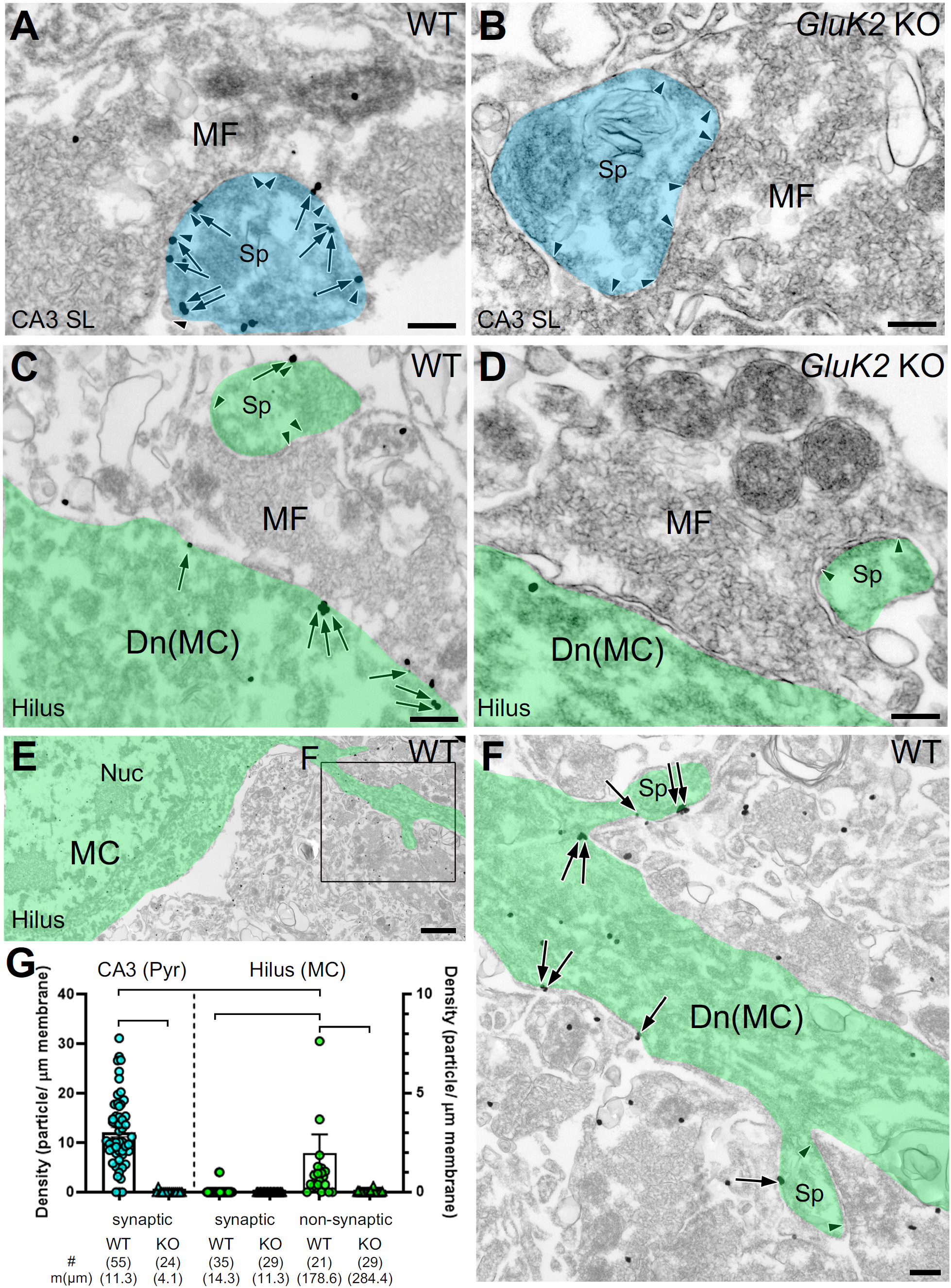
GluK2/3 in hilar MCs are enriched at non-synaptic sites. (**A,B**) A large MF terminal characteristically forms multiple asymmetric synapses (arrowheads) with TEs (blue) of CA3 pyramidal cells. In wild-type (WT) mice, metal particles for GluK2/3 (arrows) are prevalent on PSD (A). Postsynaptic labeling is absent in *GluK2* KO mouse (B). (**C-F**) MCs (green) contact with large MF terminals in the hilus of the DG. In WT mice, MF synapses on MC spines are rarely labeled for GluK2/3; occasionally, non-synaptic membrane on dendritic shaft has low but significant labeling (C). Neither synaptic nor extrasynaptic labeling is observed in *GluK2* KO mice (D). MCs occasionally have thin dendrites originating from soma (E) and contact with large MF terminal via multiple spines (F). Note that metal particle for GluK2/3 (arrows) leave postsynaptic membrane unlabeled. (**G**) Average and individual data points for the density of metal particles for GluK2/3 on CA3 pyramidal cells (*blue symbols, left axis*) and hilar MCs (*green symbols, right axis*). Note low but significant non-synaptic labeling on MCs. Edges of PSD are indicated by pairs of arrowheads. Number of measured profiles (#) and membrane length (mm) are indicated in parentheses. Dunn’s post Test. \**p* < 0.05; \*\**p* < 0.01; \*\*\**p* < 0.001. Scale bars, (A-D,F) 200 nm; (E) 1 µm.

### Activation of KARs in mossy cells by increase in ambient glutamate

Given the extracellular location of KARs in MCs we hypothesized that these receptors are activated by ambient glutamate. To test this possibility, we blocked excitatory amino acid transporters (EAATs), a manipulation that can raise extracellular glutamate and activate extrasynaptic NMDARs (Le Meur et al., 2007). We first examined the effect of the non-selective EAAT blocker DL-TBOA (100 μM) on MC holding current in presence of antagonists of AMPARs, NMDAR, GABA_A_ and GABA_B_ receptors (see Methods). Bath application of TBOA mediated a significant NBQX-sensitive inward current in MCs, suggesting that KARs could be activated by endogenous glutamate (**Fig 6A-C**) (ΔI holding MC + TBOA: 61.24 ± 18.61 pA; * p = 0.02173 one sample t Test). We next used the more selective blocker dihydrokainic acid (DHK), which selectively blocks EAAT2 (GLT-1), a glutamate transporter that accounts for ∼90% of glutamate uptake (Rose et al., 2017) and is enriched in the telencephalon including the hippocampus (Chaudhry et al., 1995). Bath application of 100 μM DHK also induced NBQX-sensitive inward currents in MCs (**Fig. 6A-C**), strongly suggesting that DHK-induced currents are mediated by activation of KARs (ΔI holding MC + DHK: 28.06 ± 7.4 pA; n = 3a/6c; n = 4a/6c). To determine that DHK-induced currents was due to activation of KARs, we repeated the experiment in *GluK2* KO mice, and found that in these mice DHK failed to induce inward currents in MCs (**Fig. 6A,C**) (ΔI holding MC + DHK *GluK2* KO: −8.5 ± 7.47; n = 3a/4c; DHK vs DHK *GluK2* KO mice: * p = 0.0105 two sample t Test). These finding suggest that increases in ambient glutamate can activate extrasynaptic MC-KARs.

**Fig 6.**
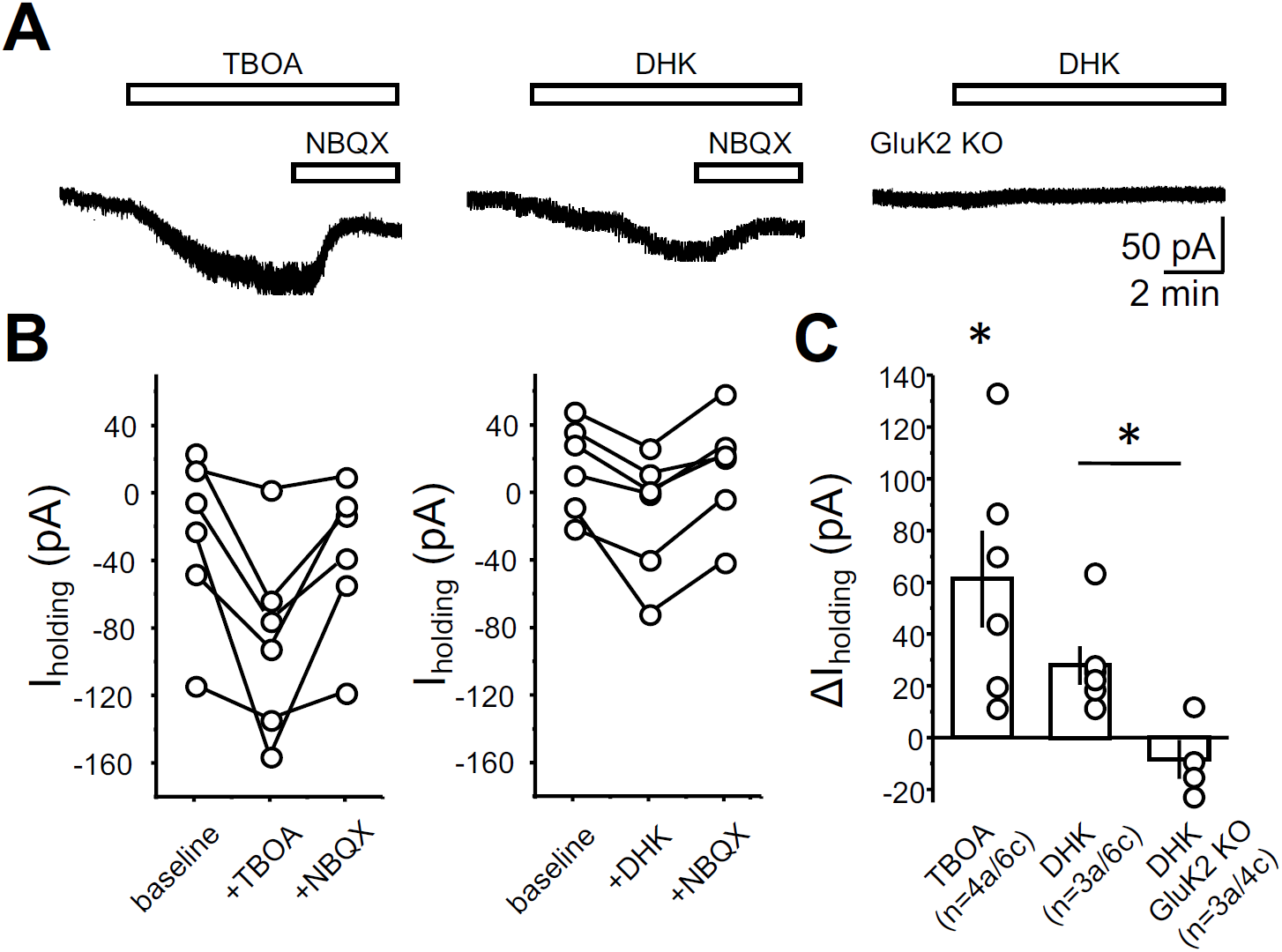
MC-KARs can be activated by endogenous glutamate. (**A**) Representative experiments showing the effect of TBOA (100 µM - *left*), dihydrokainic acid (DHK 100 µM – middle) mice and DHK in *GluK2* KO mice on MCs holding current. Recordings conditions were as in Fig.1. Activation of KARs was confirmed by application of the AMPAR/KAR antagonist NBQX (25 µM) at the end of the recording. NBQX was not applied in *GluK2* KO as no inward current was detected. (**B**) Quantification of the effect of TBOA (left) and DHK *(right*) on MCs holding current. Dots represent the average current of 2 minutes of recording taken 2 minutes before DHK/TBOA application (baseline), 2 minutes before NBQX application (+DHK/TBOA), and 2 minutes at the end of NBQX application (+NBQX). Connected dots represent the same cell. (**C**) Summary plot of the experiments shown in (B) and in the DHK *GluK2* KO dataset. (TBOA: 61.24 ± 18.61 pA; * p = 0.02173 one sample t Test; DHK: 28.06 ± 7.4 pA; n = 3a/6c; n = 4a/6c; DHK *GluK2* KO: −8.5 ± 7.47; n = 3a/4c; DHK vs DHK *GluK2* KO mice: * p = 0.0105 two sample t Test). Data are presented as mean ± S.E.M.

## DISCUSSION

In this study, we provide functional and anatomical evidence that MCs express extrasynaptic KARs whose activation in the rodent hippocampus can drive MCs. Specifically, we show that low concentrations of KA induced inward currents and action potential firing of MCs. In contrast, KA-induced currents were nearly absent in GCs, indicating that KARs have a unique pattern of expression among excitatory cells in the DG. Unexpectedly, MF activation failed to evoke a KAR-EPSCs in MCs, indicating that MF-MC synapses, unlike MF-CA3 synapses, do not normally express synaptic KARs. Our immunofluorescence and immunoelectron microscopy data confirmed that KARs in MCs are sparsely distributed at extrasynaptic sites and mainly excluded from postsynaptic compartments. Finally, blockade of the astrocytic glutamate transporter EAAT2 revealed that MC-KARs can be activated by increasing ambient glutamate.

The presence of extrasynaptic KARs has previously been suggested in hippocampal CA1 pyramidal neurons (Bureau et al., 1999), striatal medium spiny neurons (Chergui et al., 2000) and cortical layer V pyramidal neurons (Eder et al., 2003). These functional studies inferred the presence of extrasynaptic KARs given the robust effects to bath applied KAR agonist (e.g., membrane depolarization, inward current, action potential firing) with little evidence for KAR-mediated EPSCs. Of note, none of these studies provided ultrastructural evidence in support of extrasynaptic KARs. To the best of our knowledge, our immunoelectron microscopy data together with our electrophysiological characterization is the first direct evidence of a selective extrasynaptic localization of functional KARs in the mammalian brain.

Given the similarities between MF-CA3 and MF-MC synapses, the absence of KARs at MF-MC synapses is intriguing. Based on the presence of a GYKI-resistant component following MF stimulation, a previous study reported the presence of postsynaptic KARs in MCs (Hedrick et al., 2017). However, these currents were not validated in *GluK2* KO mice and showed relatively fast kinetics, which is unusual for KAR-EPSCs. Our study does not discard the possibility that KARs could be expressed at MF-MC synapses early during development (Lauri and Taira, 2011; Lerma and Marques, 2013). The molecular mechanisms that target KARs to the synapse remain unclear, but several KAR interacting proteins such as Neto 1 and 2 (Straub et al., 2011; Tomita and Castillo, 2012; Wyeth et al., 2014), N-cadherins (Coussen et al., 2002; Fievre et al., 2016) and presynaptic C1ql family proteins (Matsuda et al., 2016; Straub et al., 2016) may contribute. Consistent with data derived from a population-level transcriptomics study in the hippocampus (Cembrowski et al., 2016), we found that Neto1 signal was absent from MCs (Fig. 4), suggesting that lack of Neto1 could contribute to the low expression of KARs at the synapse (Wyeth et al., 2014). However, lack of GluK2-containing KARs also leads to reduced levels of Neto1 (Straub et al., 2011). The C-terminal domain of KARs themselves is important for the synaptic stabilization of KARs in the cerebellum (Straub et al., 2016). Additionally, phosphorylation of specific residues in the C-terminal and other intracellular regions of KARs has been implicated in the modulation of KARs function and trafficking (Wang et al., 1993; Kornreich et al., 2007; Carta et al., 2013; Zhu et al., 2014). Further studies are required to clarify the molecular mechanisms that determine the exclusion of KARs from MF-MC synapses.

Like other extrasynaptic receptors, KARs in MCs could be engaged by a rise in ambient glutamate, which can occur as a result of glutamate spillover during sustained synaptic activity (Le Meur et al., 2007; Rose et al., 2017). Glutamate could also arise from MC dendrites, as previously reported in neocortical and cerebellar neurons (Zilberter, 2000; Shin et al., 2008), and activate extrasynaptic KARs. We found that blockade of EAAT2 induces KAR-mediated inward currents likely due to the increase in extracellular glutamate. In the CA1 area of the hippocampus, activation of extrasynaptic receptors by synaptically released glutamate is limited by efficient astrocytic EAAT2 activity (Diamond and Jahr, 2000), consistent with a neuroprotective role of this transporter (Kong et al., 2012; Pajarillo et al., 2019). However, EAAT2 may saturate in other brain areas (Armbruster et al., 2016; Pinky et al., 2018). EAAT2 saturation during neuronal hyperactivity could enable the activation of extrasynaptic KARs at MCs. Alternatively, extrasynaptic KARs could be activated by glutamate released from astrocytes (Araque et al., 2014; Pal, 2015), as previously reported in CA1 GABAergic interneurons (Liu et al., 2004). There is evidence that glutamate released from astrocytes can also activate extrasynaptic NMDARs in CA1 pyramidal neurons (Fellin et al., 2004), and depolarize both hilar GABAergic interneurons and MCs (Pabst et al., 2016). These depolarizations were blocked by non-selective antagonism of all ionotropic glutamate receptors, raising the possibility that extrasynaptic KARs could be implicated. Future work is required to determine whether astrocytic processes (Gavrilov et al., 2018) could release glutamate in proximity to extrasynaptic KARs, thereby avoiding glutamate uptake by EAAT2.

Although the precise role for extrasynaptic KARs in MCs is unclear, they might detect changes in the levels of ambient glutamate and mediate tonic depolarization. *In vivo*, MCs display high level of activity compared to neighboring GCs (Danielson et al., 2017; GoodSmith et al., 2017; Senzai and Buzsaki, 2017). In standard home cage rats, MCs stain positive for the activity-dependent immediately early gene *cFos* (Duffy et al., 2013), suggesting that even at the basal level, MCs are remarkably active. It is possible that in behaving animals, where spontaneous activity is most likely higher than *in vitro*, glutamate might escape reuptake by EAATs and activate extrasynaptic KARs. During periods of particularly high activity and potential EAAT2 saturation, KARs might act as nonlinear integrators of synaptic inputs, and enhance MC output. KARs can also work in a metabotropic fashion and could potentially affect MCs excitability by suppressing the slow afterhyperpolarization (Melyan et al., 2002; Ruiz et al., 2005). Ultimately, MC-KARs could contribute to the promiscuous activity of MCs in multiple locations and environments (Danielson et al., 2017; GoodSmith et al., 2017; Senzai and Buzsaki, 2017).

Both MCs and KARs have been linked to several neurological and psychiatric disorders (Ratzliff et al., 2002; Lerma and Marques, 2013; Scharfman, 2016). Of particular relevance is the strong link between KARs and MCs with TLE. KARs have been strongly implicated in epilepsy (Crepel and Mulle, 2015; Falcon-Moya et al., 2018), and KA-induced TLE is one of the most widely used models of TLE (Levesque and Avoli, 2013; Rusina et al., 2021). The importance of KARs in KA-induced TLE is highlighted by the fact that loss of GluK2-containing KARs, strongly reduces the susceptibility to KA-induced seizures (Mulle et al., 1998). Similarly, MCs have been proposed to have a proepileptogenic role in the early phases of TLE (Ratzliff et al., 2002; Botterill et al., 2019) and to undergo prominent cell-death in both animal models of epilepsy (Blumcke et al., 2000) and in human patients (Margerison and Corsellis, 1966; Seress et al., 2009). However, the precise mechanism through which KARs and MCs are involved in TLE is still unclear. Expression of KARs in MCs strongly suggests that MCs could be a direct target of KA in KA-induced TLE. KA-induced MCs firing could contribute to hyperexcitability of the associative GC-MC-GC network and to the generation of seizures. Moreover, sustained KAR-mediated MCs firing could be a critical trigger for long-lasting forms of plasticity in the DG associative network (Hashimotodani et al., 2017), which could contribute to the prolongation of epileptic activity. A major limitation for the study of MC function in behavior is the lack of molecular tools that target MCs specifically. Thus far, manipulation of MCs activity *in vivo* relied on viral delivery of constructs under the control of promoters that are not highly specific for MCs (Jinde et al., 2012; Puighermanal et al., 2015). Establishing the precise role of MC-KARs on hippocampal function and TLE will require novel strategies such as intersectional genetics approaches (Dymecki et al., 2010; Graybuck et al., 2021) that will allow more selective targeting of MCs while sparing neighboring KAR-expressing cells such as CA3 pyramidal neurons and hilar interneurons.

## Acknowledgments

We thank the members of the Castillo lab (in particular Dr. Kaoutsar Nasrallah) for constructive feedback and for reading and commenting the manuscript. This research was supported by the National Institutes of Health: R01-NS113600, R01-MH116673 and R01MH125772 to P.E.C; Grant-in-Aid for Specially Promoted Research from Japan Society for the Promotion of Science (JSPS) (20H05628) and for Scientific Research B from Japan Society for the Promotion of Science (JSPS) (21H02589) to M.W.; JSPS KAKENHI Grants (17H0631310 and 20H03410) to M.Y.

